# A comparison of tests for quantifying sensory eye dominance

**DOI:** 10.1101/219816

**Authors:** Manuela Bossi, Lisa Hamm, Annegret Dahlmann-Noor, Steven C. Dakin

## Abstract

Clinicians rely heavily on stereoacuity to measure binocular visual function, but stereo-vision represents only one aspect of binocularity. Lab-based tests of *sensory eye dominance* (SED) are commonplace, but have not been translated to wider clinical practice. Here we compare several methods of quantifying SED in a format suitable for clinical use. We tested 30 participants with ostensibly normal vision on 8 tests. Seven tests (#1-7) were designed to quantify SED in the form of an interocular *balance-point* (BP). In tests #1-6, we estimated a *contrast-BP*, the interocular difference in contrast required for observers to be equally likely to base their judgement on either eye, whereas in test #7 we measured binocular rivalry (interocular ratio of sensory dominance duration). We compare test-retest reliability (intra-observer consistency) and test-validity (inter-observer discriminatory power) and compare BP to stereoacuity (test #8). The test that best preserved inter-observer differences in contrast balance while maintaining good test-retest reliability was a polarity judgement using superimposed opposite-contrast polarity same-identity optotypes. A reliable and valid measure of SED can be obtained rapidly (20 trials) using a simple contrast-polarity judgement. Tests that use polarity-rivalrous stimuli elicit more reliable judgments than those that do not.

**Significance Statement:** Although sensory eye dominance is central to understanding normal and disordered binocular vision, there is currently no consensus as to the best way to measure it. Here we compare several candidate measures of sensory eye dominance and conclude that a reliable measure of SED can be achieved rapidly using a judgement of stimulus contrast-polarity.

## Introduction

### Background

When combining signals from the two eyes into a coherent ‘cyclopean’ percept, observers rely (to differing extents) on one eye more than the other. Such *eye dominance* is important in a number of clinical settings. First, imbalances between the eyes in childhood can lead to severe loss of acuity in the weaker eye, a condition called amblyopia (Birch 2013). Treating amblyopia (e.g. by patching) attempts to “rebalance” vision, by improving acuity in the weaker eye. Thus the relative contribution of each eye to cyclopean vision a key measure for understanding amblyopia. This is particularly the case as there is debate as to the extent to which suppression of the weaker eye contributes to (e.g. Wong 2012, Birch 2013, Hess and Thompson 2015) or results from (e.g. Vedamurthy, Nahum et al. 2015, Kehrein, Kohnen et al. 2016, Bossi, Tailor et al. 2017) the condition, or whether this is even a reasonable dichotomy. Second, presbyopia (an age-related refractive error) can be corrected using monovision, which enhances intermediate-distance vision in the dominant eye only, and so necessitates a decision on eye dominance. Finally, loss of visual function in age-related macular-disease leads to subtle changes in patients’ reliance on their two eyes (“binocular balance”). A simple method for quantifying changes in sensory eye dominance may therefore have diagnostic value for this and other conditions (Wiecek, Lashkari et al. 2015).

A variety of methods are available to measure the relative contribution of each eye in a clinical setting. Clinical measures of eye dominance tend to be intuitive, for example assessing with which eye a patient can more easily wink (Miles 1930). Building on the notion that eye dominance is linked to motor control, comparisons of eye-alignment are still routinely used. For instance, in the *Miles test* and *Porta test*, near and distant targets are aligned binocularly, and then each eye is opened and closed in succession to reveal which eye gives the more accurate estimate of position. Such tests of ‘sighting’ dominance are binary (left or right) and tend to elicit variable results (Johansson, Seimyr et al. 2015), that are dependent on specific test and viewing conditions (Rice, Leske et al. 2008).

Sensory eye dominance (SED), on the other hand, refers to the relative *perceptual* contribution of each eye to the cyclopean percept under dichoptic conditions (Ooi and He 2001). In the clinic, *Worth’s four dot test* produces a qualitative estimate of SED. This task has the observer wear red-green anaglyph glasses and report the perceived number and colour of 4 illuminated circles (1 red, 2 green and 1 white). The only *quantitative* direct clinical test of SED is the *neutral-density filter bar* (a version using red filters is the *Sbisa bar*), that quantifies the minimum filter-density that must be placed over one eye to induce the use of the other eye (McCormick, Bhola et al. 2002). However, such tests can be challenging for children which likely contributes to their only moderate test-retest reliability (Crawford and Griffiths 2015, Piano and Newsham 2015) and their infrequent use for paediatric screening (Tailor, Balduzzi et al. 2014).

Neither sighting-dominance nor sensory dominance assess the functional advantage of having two eyes. When each eye makes a relatively equal contribution, observers experience binocular summation (increased contrast sensitivity when using both eyes; Baker, Meese et al. 2007, Baker, Meese et al. 2007, Baker, Meese et al. 2008) and stereopsis (the perception of three-dimensional structure based on retinal disparity; Cumming and DeAngelis 2001, O’Connor, Birch et al. 2010). Although there is no standard clinical test of summation, stereoacuity is the most widely accepted clinical measure of binocular vision. Stereoacuity tests quantify the minimum retinal disparity supporting reliable discrimination of surface-depth (Julesz 1971). Popular variants include the *TNO*, the *Randot* test (Birch, Williams et al. 2008) and the *Frisby* test (Frisby, Davis et al. 1996). Stereoacuity tests are simple to administer and can easily be explained to children. However, they have drawbacks; some contain monocular cues (Fricke and Siderov 1997), they can be difficult to administer in the presence of strabismus (McKee, Levi et al. 2003) and they are not necessarily indicative of clinical conditions (since 1-14% of the general population are stereo-blind; Bosten, Goodbourn et al. 2015) which limits their utility for vision screening (Cotter, Cyert et al. 2015). For example, despite being stereo-blind patients with strabismic amblyopia can demonstrate normal binocular summation when balancing the visibility of dichoptic stimuli (Baker, Meese et al. 2007).

Outside the clinic, where stereoacuity is the foundation of binocular assessment, vision researchers have developed diverse, innovative and quantitative measures of sensory eye dominance. Rather than directly asking observers to make a judgement about contrast similarity (as required for e.g. the Sbisa bar), psychometric assessment tends to rely on forced-choice methods (for a study that linked traditional clinical methods of SED to a common psychophysical measure see Li, Lam et al. 2010). Pairing robust psychophysical judgements with carefully-designed stimuli allows assessment of the relative contribution of each eye to visual processing in different stages of the cortical hierarchy.

Here we consider SED tests first according to whether they do or do not preclude a coherent cyclopean percept (i.e. are intrinsically *rivalrous*). One test which poses no cyclopean conflict presents observers with dichoptic sine-wave gratings that differ only in phase and contrast (Figure 1, test #4). Observers are asked to indicate the location of the middle dark stripe of the phase-shifted grating that results from binocular summation (Ding and Sperling 2006, Huang, Zhou et al. 2009, Huang, Zhou et al. 2010, Zhou, Thompson et al. 2013, Kwon, Lu et al. 2014). Reported position allows one to determine the interocular contrast difference that supports equal contribution from each eye, and it is this contrast difference that quantifies “binocular balance”. Another test embeds contrast manipulation within a global processing task (Mansouri, Thompson et al. 2008, Li, Lam et al. 2010, Black, Thompson et al. 2011, Li, Hess et al. 2013, Li, Hess et al. 2013, Li, Thompson et al. 2013, Hamm, Chen et al. 2017). In motion coherence paradigms (Li, Lam et al. 2010), the observers are required to discriminate the direction of a pattern of moving dots, comprised of signal-dots (moving in one direction) and noise-dots (moving in random directions). The most commonly used version of this approach for quantifying SED requires two phases: (1) determining the proportion of signal to noise dots supporting reliable direction discrimination and (2) determining the contrast of the signal-dots required to maintain performance under dichoptic conditions (Black, Thompson et al. 2011). This paradigm has also been used with global orientation tasks (Zhou, Huang et al. 2013). In all such tests, observers perceive a coherent cyclopean percept and the relative contrast supporting optimum performance is termed the interocular ‘balance point’ and quantifies SED.

**Figure 1.**
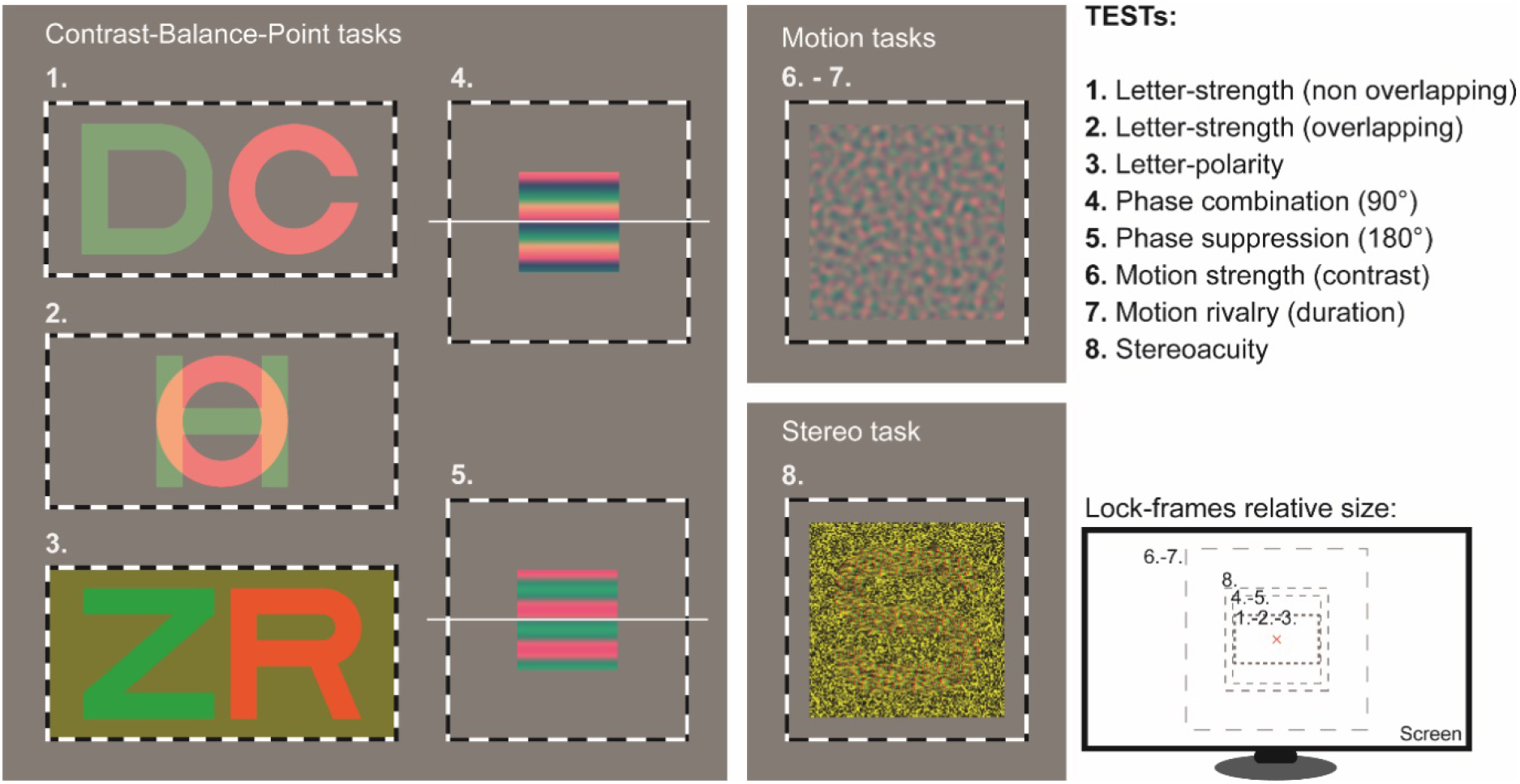
(Grey panels) Examples of stimuli used in the eight tests (for the purpose of demonstration, stimuli are presented in red-green anaglyph versions). (Inset, lower right) Relative sizes of stimuli.

A second category of tests use rivalrous stimuli to quantify SED. When different monocular images fall on corresponding retinal locations of the two eyes this can lead either to an experience of *binocular rivalry* (an alternation of percept between the stimuli presented to the two eyes; Wheatstone 1838), or to *diplopia* (double vision). The nature and extent of conflict between the information from the two eyes is a measure of the degree of SED. Bossi *et al* (2017) quantified this using dichoptic stimuli where each eye was presented with opposite contrast-polarity versions of the same symbol (Figure 1, test #3), whereas Kwon et al presented observers with spatially overlapping rivalrous letter pairs of differing contrast (Figure 1, test #2; Kwon, Wiecek et al. 2015). In both cases, the objective was to measure the ‘contrast balance point’, or the contrast mixture required for observers’ percept to be equally likely to be driven by either eye’s view. This balance point (like the non-rivalrous tests) quantifies SED.

Although all tests described use centrally presented stimuli, SED has been measured at multiple locations in the visual field. Xu et al mapped SED over 17 locations (at the fovea, 2 and 4 deg. eccentricity) and report gradual variation across the field (Xu, He et al. 2011). Hess and colleagues have measured contrast balance in patients with strabismus and report that patients exhibited stronger suppression across the field than controls (Babu, Clavagnier et al. 2017). Development of this, and many of the tests described, has been driven by research exploring the role of interocular suppression in amblyopia (Baker, Meese et al. 2008, Mansouri, Thompson et al. 2008, Huang, Zhou et al. 2009). Several studies have examined the use of training (or *perceptual learning*) to shift balance towards the amblyopic eye as a treatment for amblyopia (Ooi, Su et al. 2013, Birch, Li et al. 2015). Although the benefits of such training for acuity in the amblyopic eye are clear, it is less clear if these benefits result from a shift in SED (Vedamurthy, Nahum et al. 2015, Bossi, Tailor et al. 2017) as has been reported(Black, Hess et al. 2012). Although the functional significance of changes in SED is debatable, it is the case that such changes can arise from short term manipulation of binocular sensory experience (e.g. through patching) in both amblyopes (Zhou, Thompson et al. 2013) and controls (Zhou, Clavagnier et al. 2013, Zhou, Reynaud et al. 2014).

There have been several efforts to translate lab-based measure of SED to the clinic. Typically, research groups have adapted their own method for clinical use - making tests more convenient/shorter/simpler (e.g. Black, Thompson et al. 2011, Kwon, Lu et al. 2014). Here we compared a variety of dichoptic stimuli using a common psychophysical protocol. We assessed resulting SED scores based on 1) *Reliability* (consistency between two measurements made on the same participant), and 2) *Validity* (the ability of the test to capture individual differences).

We selected (or created) tests suitable for a single, rapid psychophysical protocol (effectively precluding 2-stage paradigms and tasks involving multiple stimulus locations). We ensured tasks probed a range of visual processes; from low-level (e.g. phase; Ding and Sperling 2006), to high-level processing of spatial form (e.g. letter identity; Kwon, Wiecek et al. 2015), as well as motion perception. We also set out to quantify the impact of rivalry on our measures, focusing on stimuli involving no-rivalry (Ding and Sperling 2006), contrast-polarity rivalry (Bossi, Tailor et al. 2017) and spatial-form rivalry (Kwon, Wiecek et al. 2015).

## Methods

### Participants

We recruited thirty adult participants (18 female; 22-55yrs old) through the University of Auckland-Optometry clinic via email and poster advertisements. Based on pre-test screening, none had diagnosed disorders of vision and all wore habitual correction as necessary. Participants’ acuity and refraction were determined from clinical notes or using an auto-refractor and 3m Sloan letter logMAR chart. All participants had visual acuity <0.2 logMAR in both eyes except ID24 whose left eye acuity was 0.2 logMAR. The maximum inter-ocular difference in acuity in our group was 0.1 logMAR.

Three of the observers are authors; others were not informed as to the purpose of this study. Experimental protocols complied with the Declaration of Helsinki and were approved by the University of Auckland Human Research Ethics Committee.

### Apparatus

Stimuli were presented on a linearised LG 21” LCD 3D monitor, with a 1920×1080 pixel resolution operating at 120Hz. Stimuli were generated in Matlab (Mathworks Ltd) using the Psychophysics Toolbox (Brainard, 1997; Pelli, 1997; Kleiner et al, 2007). During testing, stimuli were viewed through wireless LCD shutter glasses (nVidia Corp., Santa Clara, CA) allowing independent control of the image presented to each eye (effective framerate was 60Hz per eye). Shutter glasses were worn over optical correction when necessary. A Minolta LS110 photometer (Konica-Minolta Ltd) was used to calibrate the monitor-luminance, using measurements made through the shutter glasses. Participants viewed the screen at a viewing distance of 100 cm to produce a pixel density of 55.9 pixel/deg.

### Stimuli

Red-green anaglyph versions of the stimuli are depicted in Figure 1, and described in Table 1. Stimuli appeared against a mid-grey background, and were surrounded by a “vergence-lock” frame (visible to both eyes) made of black and white alternating bars; this promoted fusion.

**Table 1.** Summary of eight tests. BP= Balance Point (either contrast-BP in tests #1-6 or time-ratio BP in test #7).

| Test # | Test name | Symbol | Cue | Rivalry? (type) | Question | Measure | Primary reference |
| --- | --- | --- | --- | --- | --- | --- | --- |
| 1 | Letter-strength (non-overlapping) | NH | form | ✗ | Which letter appears stronger? | Contrast-BP | Kwon, Wiecek et al. (2015) |
| 2 | Letter-strength (overlapping) | NH | form | ✓ (form) | Which letter appears stronger? | Contrast-BP | Kwon, Wiecek et al. (2015) |
| 3 | Letter-polarity | ZK | form | ✓ (polarity) | Which letter appears whiter? | Contrast-BP | Bossi, Tailor et al. (2017) |
| 4 | Phase combination | Horizontal bars | position | ✗ | Does middle dark bar fall above or below horizontal ref. line? | Contrast-BP | Ding and Sperling (2006) |
| 5 | Phase suppression | Horizontal bars | position | ✓ (polarity) | Does middle dark bar fall above or below horizontal ref. line? | Contrast-BP | Ding and Sperling (2006) |
| 6 | Motion strength (contrast) | Vertical arrows | motion | ✓ (motion) | Does the pattern move up or down? | Contrast-BP | Black, Thompson et al. (2011) |
| 7 | Motion rivalry (duration) | Vertical arrows | motion | ✓ (motion) | Hold ↑ key when pattern moves ↑ & ↓ key when it moves ↓ | Time-ratio BP | Dieter, Sy et al. (2016) |
| 8 | Stereoacuity | 3D letter S | form | ✗ | Identify the letter you see | Threshold disparity | Frisby, Davis et al. (1996) |

All tests involved presenting a pair of images independently to each eye. **Test #1-2**. Stimuli comprised a pair of different letters, selected from 10 ETDRS Sloan letters (Ferris, Kassoff et al. 1982). The width and height of letters was 2.75° and the letters were the same (positive) contrast polarity. In test #1 letters were spatially separated (minimising rivalry) whereas in test #2 they were superimposed, promoting rivalry between the two different letters. **Test #3** Letter font, size and separation were identical to test #1. However, each of the two letters was comprised of superimposed pair of opposite contrast polarity letters (with the same identity) presented separately to the two eyes. In one stimulus-letter the light component went to the left eye, the dark component to the right, and in the other stimulus-letter, vice-versa. This led to strong rivalry, driven by the conflicting contrast-polarity of the components of each letter. **Tests #4-5**. Stimuli were similar to Ding & Sperling (2006) and were comprised of two horizontal sine-wave gratings with the same spatial frequency (SF; 0.52 c/deg.), presented within a 2.94° square window. The two gratings were presented separately to each eye and had phases of ±45°(test #4, 90° phase difference, Figures 1–4) or ±90° (test #5, 180° phase difference, Figures 1–5), where a 0° phase grating would present a horizontal dark bar centrally positioned (on the y-axis) bisecting the square window. On a given trial the positive-phase grating was assigned randomly to one eye, the negative-phase grating to the other. In the 90° condition (test #4) addition of gratings led to a phase shifted cyclopean percept without rivalry, whereas in the 180° condition gratings resulted in rivalry (driven by a polarity conflict similar to test #3). **Tests #6-7**. Stimuli were comprised of two 13.75° square filtered-noise patterns drifting in opposite directions (upwards or downwards). Stimuli were generated by filtering Gaussian random noise with a log Gabor filter which passed all orientations but a range of SFs (filter had a log-Gaussian profile: mean SF of 6 c/deg., bandwidth 0.5 octaves). **Test #8**. Stimuli were stereograms - stereo-defined Sloan letters (5.5° square) - defined by interocular horizontal shifts of the pixels within carrier images (435 X 435 black-white pixel arrays which each patch subtending 7.8°). Stimuli were shifted with sub-pixel accuracy using bilinear interpolation.

### Procedure

We ran participants through eight tests, twice (to allow us to estimate test-retest reliability). Tests were administered in a fixed order (#1-8). Tests were ordered in this way because we sought to maximise the participant’s engagement and to minimise the impact of fatigue. We found the letter tasks (#1-3) to be the most intuitive for participants to grasp. Test #1 is non-rivalrous and allows the participant to see both letters at once (making the stimulus easy for them to understand), test #2 extends test #1 by introducing form-rivalry, and test #3 changes the rivalry to polarity-rivalry. Tests #4 and 5 are the phase tasks which can be more difficult for some to grasp but practice on tests #1-3 helps. Again, we start with a non-rivalrous test (#4) and introduce rivalry (#5). Test #6 is the last and most challenging test because of the “patchy” rivalrous appearance of stimuli. Test #7 is similar except the components are of a fixed contrast and participant must use the keyboard themselves to signal changes (the only test that requires this). Finally, we decided to leave the unique stereopsis test #8 to the end (of each sequence).

Participants first performed all tests once (run 1) and then repeated the whole test-sequence (run 2). In all tests except #7 (motion rivalry) stimuli appeared for a maximum of 4.5 s, after which the display switched to showing the response-choices for a maximum of 10s. Observers could respond before the display of response options, which triggered presentation of the next trial. Participants signalled their response verbally, which the experimenter recorded using the computer keyboard. Note that no feedback was provided in any test (since we are estimating bias in all but test #8). Each test took approximately 90s to administer. The experimenter instructed the participant before each test as highlighted in Table 1. Each test took no more than 1 min for participants to perform and was separated from the upcoming test by a 1-2 min break. During this time the experimenter described the next test and participants were free to look around the room. This schedule minimised any impact of testing on subsequent tests (e.g. because of adaptation). Total duration of the experiment - incorporating 8 tests x 2 repeats, instructions and breaks - was around 50 min. Tests #1-7 measured a Balance Point (BP) from 0 (participant used their left eye only), to 1 (right eye only). For tests #1-6 BP was based on contrast, and for task #7 it was based on a ratio of time spent experiencing the stimuli presented to each eye.

**Tests #1-6** quantified the ratio between the contrast-level applied to each dichoptic-image that leads to either of the two images (called A, B below) being equally likely to be chosen. This contrast-BP was determined in 20 trials using an adaptive staircase algorithm (QUEST; Watson and Pelli 1983) which, for a stimulus comprised of the mixture of images *A* and B, converged on a threshold (α) producing 50% identification of (tests #1,2,4-6) image *A* or (test #3) the “whiter” stimulus. The values QUEST produced were clamped in the range 0.0-1.0 (where 0.0 is exclusive presentation of the cue to the left eye, 0.50 equal presentation to left and right eye and 1.0 exclusive presentation to the right eye). QUEST’s guess-rate (γ) was set to 0.5 (2 AFC), the lapse rate (λ) to 0.01 and the slope-estimate (β) to 3.5. The initial guess for threshold or contrast-BP was 0.50 (i.e. balance) with an associated standard deviation of 0.7. For tests #1, 2, 4, 5 and 6, QUEST set the contrast (C) of the right eye component (*C*_right_) and *C*_left_ was set to 1-*C*_right_, for test #3 contrast manipulation is described in the next section. Note that QUEST values were determined from a Monte Carlo simulation on a population of ideal observers whose simulated-balances spanned the range 0.05-0.95 and whose other psychometric characteristics (β, λ) were taken from Kwon, Wiecek et al. (2015). **Test #7** consisted of 1 minute of exposure to rivalrous motion. Either upward or downward motion was randomly assigned to the left and right eyes for the initial 30s test period and then direction was switched across eyes for the remaining 30s (to counter-balance any bias for a stimulus in a given direction rather than from a given eye). During stimulus presentation, observers pressed and held either the U (“up”) or D (“down”) button on the computer keyboard to indicate their dominant percept. In case of uncertainty, e.g. due to a “patchy” percept, the participant was instructed to report the more dominant direction. **Test #8** estimated a stereo-threshold (not bias) for letter-identification, using a 20-trial QUEST staircase. QUEST’s guess-rate (γ) was set to 0.1 (10 AFC), the lapse rate (λ) to 0.01 and the slope (β) to 3.5. The initial guess for threshold was set to 3’20”.

**Details of contrast manipulation for test #3** For test #3, the QUEST value set the luminance of the components of the opposite contrast polarity left/right-eye letters forming each of the two stimulus letters. First, the highest increment was randomly assigned to the stimulus-letter *A* or B. Then, the luminance of the light component-letter in one stimulus-letter and the dark component-letter in the other stimulus-letter were set to be equal increments and decrements (range ±50cd/m^2^; example in Figure 2 of Bossi, Tailor et al. 2017). Thus, for a background luminance of 50 cd/m^2^ if QUEST produced a value of 0.8 then the light component letter in, for example, stimulus letter A was 50+0.8*50=90 cd/m^2^ and so the dark component letter in stimulus letter B was 50-0.8*50=10 cd/m^2^. The dark component letter in stimulus letter A was 50-(1.0-0.8)*50=40 cd/m^2^ and the light component letter in the stimulus letter B was 50+(1.0-0.8)*50=60 cd/m^2^. In the extreme case of QUEST producing the limit value of 0.0, then the light component letter in, for example, stimulus letter A was 50+0.0*50=50 cd/m^2^ and so the dark component letter in stimulus letter B was also 50-0.0*50=50 cd/m^2^. The dark component letter in stimulus letter A was 50-1.0*50=0 cd/m^2^ and the light component letter in the stimulus letter B was 50+1.0 *50=100cd/m^2^. Conversely, when QUEST converged to 1.0, the dark component of A and the light component of B were matched (at 50 cd/m^2^) while A had a light component at 100 cd/m^2^ and B a dark component at 0 cd/m^2^.

**Figure 2.**
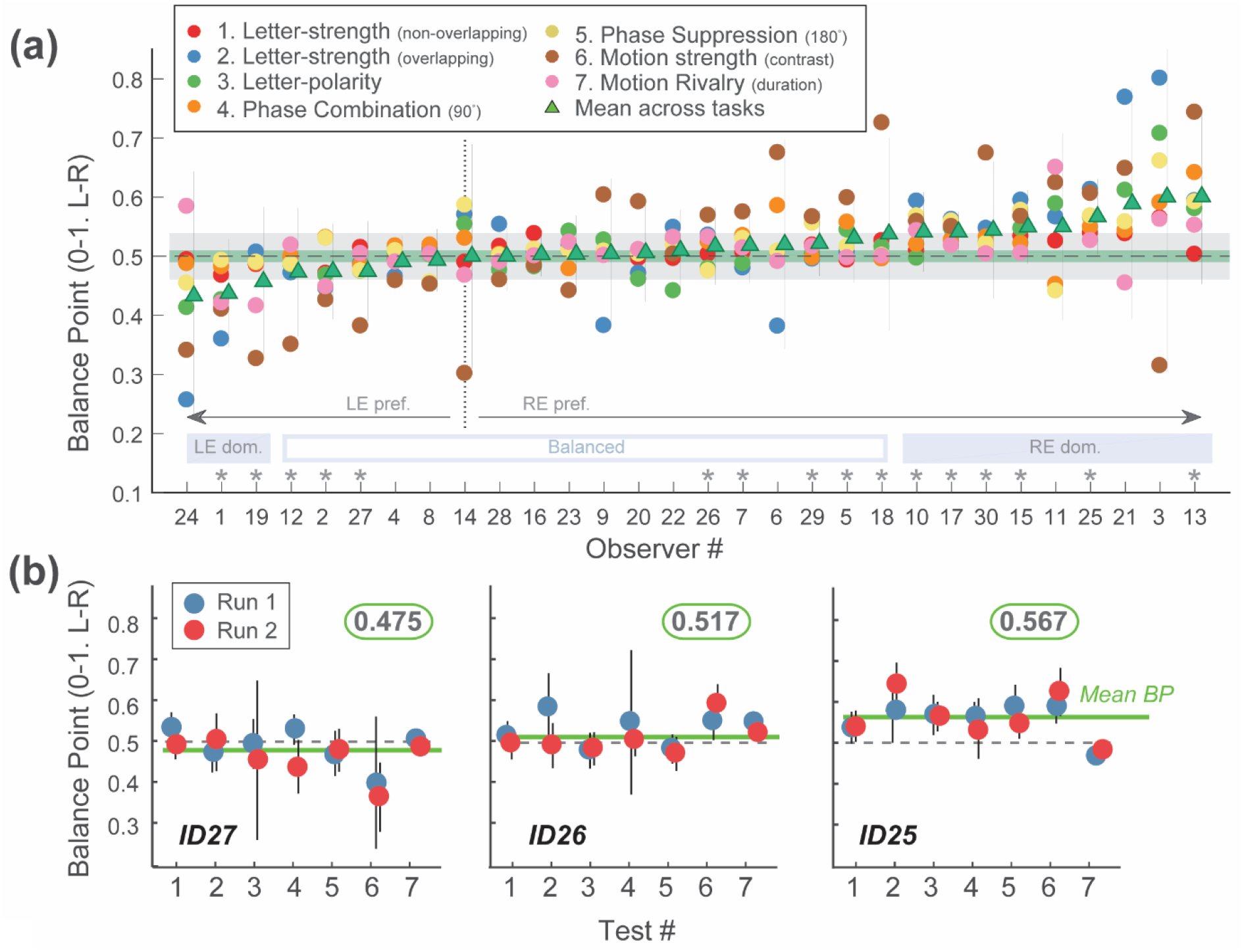
**(a)** Estimated balance point (BP) from tests #1-7, for all participants. On the y-axis, 0.0 and 1.0 represents complete reliance on the left and right eye, respectively and 0.50 equal reliance on each eye (‘Balanced’). Each coloured dot displays the mean of run 1 and run 2 for a particular BP test, according to the legend. Mean-BP for each participant is indicated by a green triangle (error bars are 95% confidence intervals). The x-axis is ordered by mean-BP. The horizontal shaded green and grey regions capture 0.5±0.5σ_all_ and 0.5±2σ_all_, respectively, where σ_a_n (=0.021) is the standard deviation of the difference in BP between runs across all participants and tests. Eye preferences is whether the green triangle falls to the left or right of the vertical dotted line (LE or RE ‘pref.’). Eye dominance is whether the green triangle falls outside the grey shaded region (LE or RE ‘dom.’). ‘Task-independent eye-dominance is whether the green triangle falls outside the green shaded region and no coloured dot falls beyond the opposite grey shaded region (summarised with an ‘*’ over each relevant observer #). **(b)** Illustrative results from three participants, chose from the left, central and right batches of IDs on the abscissa in 2a. Blue and red symbols show data from the first and second run respectively, and error bars are 95% confidence intervals on estimates. Mean BP across tasks (green line) is the green-boxed figure in each panel.

### Analysis

For tests #1-6, BP was derived by fitting the binary response data from the 20 trials of a single QUEST-controlled run with a cumulative normal function using the Palamedes toolbox (Prins and Kingdom 2009). Further fitting was performed in order to quantify confidence in the BP estimate. In order to do this, we pooled stimulus-levels (and their associated responses) into four bins, assumed binomially distributed errors at each level and bootstrapped fits to these binned data. Although this is not an exact estimate of error (since we cannot bootstrap the continuous data used to derive the contrast-BP) it is nonetheless a useful indicator of confidence in these estimates.

For test #7 we quantified BP as the proportion of left-eye dominance, using the proportion of frames (within a given trial) when the participants’ response was consistent with their relying on the left-eye-view during exposure to rivalrous stimuli. We excluded responses occurring within 4.2 s (i.e. 500 frames) of the beginning, and the middle (when directions switched between eyes) of the 60s sequence. Responses composed of simultaneous keypresses were excluded from analysis.

We first describe BP for each individual and each test by averaging run 1 and 2 together, and also calculate the mean BP for each participant across all tests. To estimate test-reliability we calculated Bland Altman 95% limits of agreement, or Coefficient of Repeatability (CoR - lower values indicate higher repeatability; Vaz, Falkmer et al. 2013) as well as the Intraclass Correlation Coefficient (ICC - higher values indicate higher repeatability), based on a mean rating (k=2), absolute agreement, 1-way random-effects model (McGraw and Wong 1996). We examine how CoR and ICCs are influenced by the number of trials run (by re-calculating BP with Palamedes refits for fewer trials), and investigate the reliability of tests by comparing measures to one another. We note that highly reliable outcomes can arise from a measure which has low test-validity, being insensitive to differences in SED between individual (in other words, a test which always elicited the same outcome would be highly reliable, but not very useful). The absence of a gold standard measure of SED means we cannot directly assess which test is the most accurate by comparing it to such a standard. We therefore relied on Mean Average Precision (MAP) and Fractional Rank Precision (FRP) to incorporate both reliability *and* validity (Dorr, Elze et al. 2017). FRP employs an information retrieval approach and evaluates a test by quantifying how identifiable a participant is from their set of test-scores. Finally, we use regression to compare SED data to stereoacuity.

## Results

Tests #1-7 estimated observers Balance-Point (BP): #1-6 estimated observers’ contrast Balance-Point, #7 quantified rivalry; test #8 measured stereoacuity. We assessed tests according to their 1) *reliability* (consistency between two measurements of the same test made on the same participant) and 2) *validity* (the ability of the test to capture individual differences amongst participants). Reliability was quantified using the Bland Altman Coefficient of Repeatability (CoR; the variance of the difference between runs 1 and run 2). The other metrics in table 2 provide statistical estimates of both reliability and validity.

**Table 2.** Summary statistics for performance of the eight tests. *Indicates best performance (across tests #1-7) for a given metric. *ICC*: intraclass correlation, F: ANOVA *F* statistic, *CoR*: coefficient of repeatability (lower is better), *MAP*: mean average precision (higher is better), *FRP*: fractional rank precision (higher is better; Dorr, Elze et al. 2017). Note CoR measures for Stereo are unique in being expressed in arc sec.

|  | *ICC* | *F* (*p*) | *CoR* | *MAP* | *FRP* |
| --- | --- | --- | --- | --- | --- |
| 1. Letter-strength (non-overlapping) | 0.60 | 4.04 (<0.001) | **0.058\*** | 0.17 | 0.64 |
| 2. Letter-strength (overlapping) | 0.73 | 6.34 (<0.001) | 0.22 | 0.20 | 0.65 |
| 3. Letter-polarity (overlapping) | **0.80\*** | **9.15\* (<0.001)** | 0.11 | **0.30\*** | **0.78\*** |
| 4. Phase-comb (90°) | 0.23 | 1.60 (=0.10) | 0.17 | 0.17 | 0.59 |
| 5. Phase-suppression (180°) | 0.63 | 4.39 (<0.001) | 0.12 | 0.20 | 0.71 |
| 6. Motion strength (contrast) | 0.32 | 1.92 (=0.040) | 0.49 | 0.26 | 0.75 |
| 7. Motion rivalry (duration) | 0.20 | 1.50 (=0.14) | 0.21 | 0.14 | 0.54 |
| 8. Stereo | 0.39 | 2.29 (=0.014) | 40.20 | 0.22 | 0.69 |

*Are observers more reliant on their left or right eye?* Figure 2a plots BPs estimated from two runs of tests #1-7 for all participants. Green symbols show the mean BP calculated by averaging contrast BPs (across tests #1-6) and proportion of percepts determined by the left eye (test #7). Mean BP estimates cluster around 0.50, indicating binocular balance, as one would expect for observers with ostensibly normal binocular vision. Note, however, the general shift towards reliance on the right eye (mean BP >0.5). To explore this, we consider three criteria for categorising whether a participant was reliant on one eye over the other (each summarised at the bottom of Figure 2a; ‘LE’=left eye, RE=‘right eye’). Criteria 2 and 3 are based in part on test reliability: σ_all_ =0.021, the standard deviation of the difference in performance between runs across all participants and balance tests.

1. *“Eye preference”*: mean BP < or > 0.50 (i.e. in Figure 2a: ‘LE pref.’ or ‘RE pref.’, respectively). By this criterion, most participants (21) showed a preference for the right-eye, eight participants showed a preference for the left-eye, and one participant (ID14) was balanced.
2. *“Eye dominance”*: mean BP <0.50-2σ_all_ or >0.50+2σ_all_ (i.e. in Figure 2a: ‘LE dom.’ or ‘RE dom.’, respectively). By this more conservative criterion, most participants (18) were balanced, nine relied more on their right eye and three relied more on their left eye.
3. *“Task-independent” eye-dominance*: mean BP < 0.5-0.5σ_all_ (favouring LE) or > 0.5+0.5σ_all_ (favouring RE) with the result of no single test indicating dominance of the opposite eye (based on 0.5±2σ_all_) to the mean BP. Observers fulfilling this criterion are marked with a ‘*’ over their ID number. By this standard, 14 participants were balanced, 11 showed a consistent reliance on the RE and 5 participants showed a consistent reliance on the LE.

Tests #1, 3, 4, 5 and 7 elicited BPs more similar to one another (and to the ovrall mean balance) compared to tests #2 and 6 (Figure 2a). Figure 2b plots a sample of data from three participants performing two runs of the seven balance tests. The dashed line denotes perfect binocular balance (BP=0.50). The green line indicates the mean estimated balance-level across the seven tests (with exact values provided in the green box above each plot). We found a generally high level of agreement across runs of the same test: the absolute difference was 0.048 ± 0.025 between participants (averaging, for each participant, the difference across tests) and 0.048 ± 0.028 between tests (averaging, for each test, the difference across participants). We examined the consistency of balance estimates using a correlation analysis. Specifically, for each task we correlated the series of contrast-BPs (across all observers) for one task (averaged across runs 1 and 2) with the series of contrast-BPs (across all observers) averaged across both runs and all other tasks. We observe consistency of contrast-balance estimates made using different tests as indicated by the mean correlation coefficient (mean R= 0.56, σ=0.10, p=0.0012) obtained by averaging the R from each task.

*Which is the “best” test?* We assessed ‘best’ in terms of (a) how reliable the tests are (across two runs) and (b) how well they capture individual variation in SED across our group. In other words, the best test minimizes the range of measures across runs (narrow range on the y-axis), and assuming that SED varies within the population tested, maximizes the range of measures across observers (wide range on the x-axis). To visualise this, Figure 3 shows Bland Altman plots of data from the seven balance tests as well as test #8: stereoacuity (n.b. data are plot on different axes to other sub-plots). It is clear that some tests such as #6: motion strength, lead to a wide range of balance estimates across observers but also to high variability of balance-estimates across runs (see also Figure 2). Conversely, tests such as letter-strength (Figure 3, task #1) elicit both a narrower range of balance estimates and much lower variability across runs. However, the high degree of repeatability of this test arises from it yielding a BP estimate of 0.50 for almost all participants, suggesting it is unable to differentiate subtle difference in SED (i.e. it has poor test-validity). We note that it is the closest test we have to the *Sbisa bar*, in that there is not spatially overlapping information and participants are required to make a judgment of contrast.

**Figure 3.**
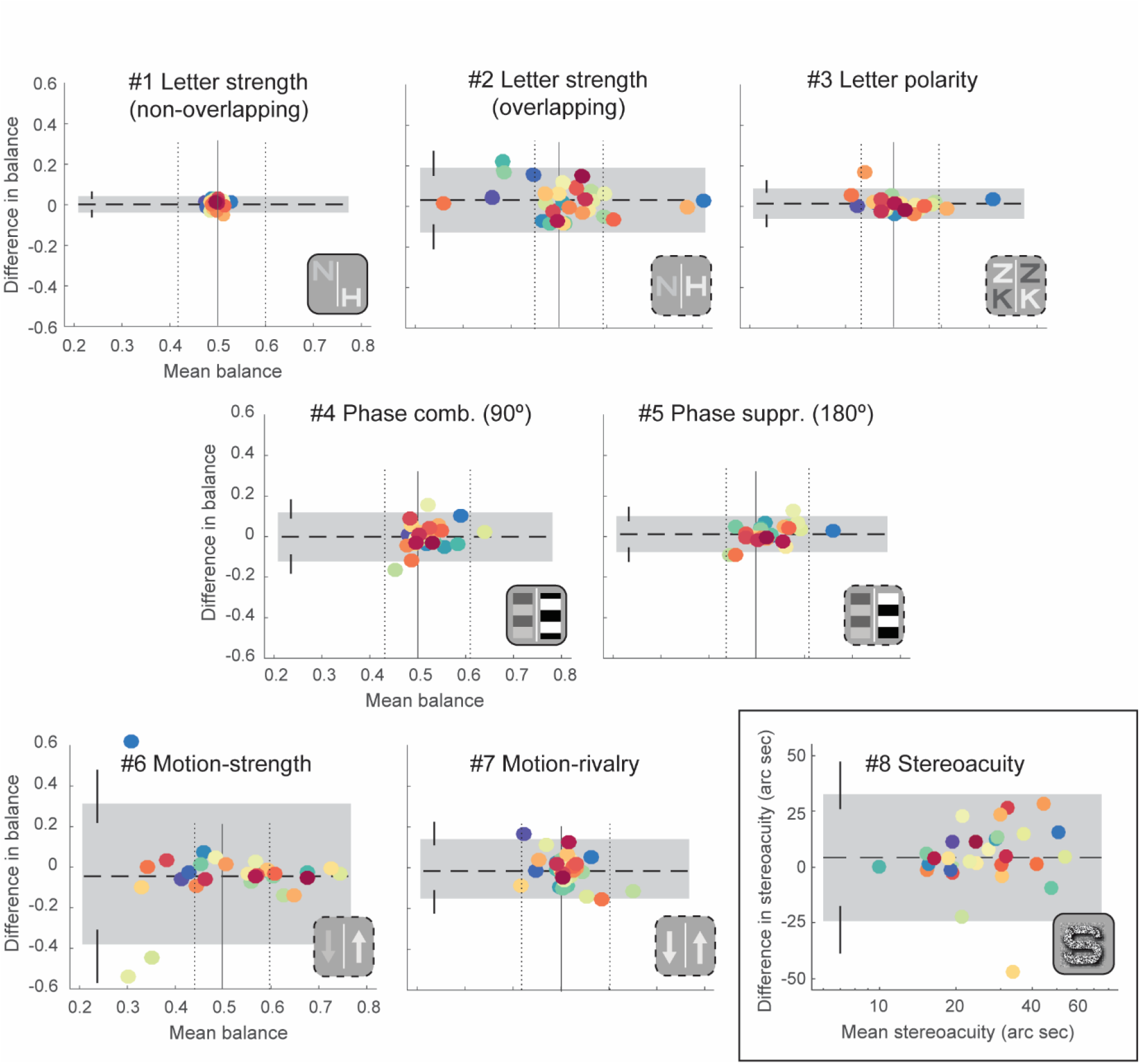
Bland-Altman plots of results from the eight tests. (Subplots #1-6) Mean contrast balance across runs is plotted against the difference in contrast balance. (Subplot #7) Relative time spent in each possible rivalrous state plot against the different in this measure across runs. (Subplot #8) Mean stereoacuity plot against the difference in stereoacuity across runs. Shaded regions denote the 95% confidence intervals across estimates with error bars on these estimates calculated using the procedure given by Carkeet (2015). In each subplot (#: 1 to 7), the solid vertical line indicates SED=0.50 (no eye-dominance) while the two flanked, dotted lines indicate 0.50±1.96*σ (between participants) of the mean balance-points averaged across all the other tests. Note high repeatability in (#1) but restricted range of estimated balance across participants. The motion-strength test (#6), in contrast, produced a broader range of estimated balance but at the expense of repeatability. (#3) The letter-polarity test combines high repeatability with a broad range of estimated contrast-balance.

A variety of statistics quantify this and a selection are given in Table 2. Test #3 (letter-polarity) maximises intra class correlation (ICC), F, mean average precision (MAP), and fractional rank precision (FRP). The Coefficient of Reliability (CoR) is lowest for test #1 - the letter-strength (non-overlapping) task. However, as indicated above, it would appear that this reliability comes at the expense of failing to capture individual variation in SED.

Based on ICCs and corresponding F values, tests #2 and #3 appear particularly useful. However, ICCs are driven by values at the ends of the measured range, making them susceptible to outliers. In our data set, participant ID3 reported quite unbalanced but reliable scores on tests #2, 3 and 5. This individual may be an outlier, inflating ICC scores for these tests. MAP and the related FRP estimates circumvent this issue by scoring on rank rather than absolute value. This difference in method impacts test #6 the most, as it has a poor ICC (and F), but fair MAP and FRR values.

Figure 4 shows the variation of the CoR and ICC over run-length. Here we analysed from 25 to 100% of data (i.e. either trials 5-20, for tests #1-6 and 8 or from 4.2-30s and 34.2-60s in test #7). For each test (colour coding given in the legend), CoR-coefficient of repeatability (Figure 4a) and ICC-interclass correlation (Figure 4b) indexes are obtained by comparing first and second run across all observers. Predictably, there is better agreement towards the end of runs, as evidenced by the descending (Figure 4a) or ascending (Figure 4b) trend of the plotted lines. Figure 4c visually represents CoR (x-axis; lower is better) against ICC (y-axis; higher is better) indexes for data from the whole run for all of the tasks tested. Note the clustering of results. Results from the two motion- and the non-rivalrous phase-combination tests were substantially poorer than the other tests.

**Figure 4.**
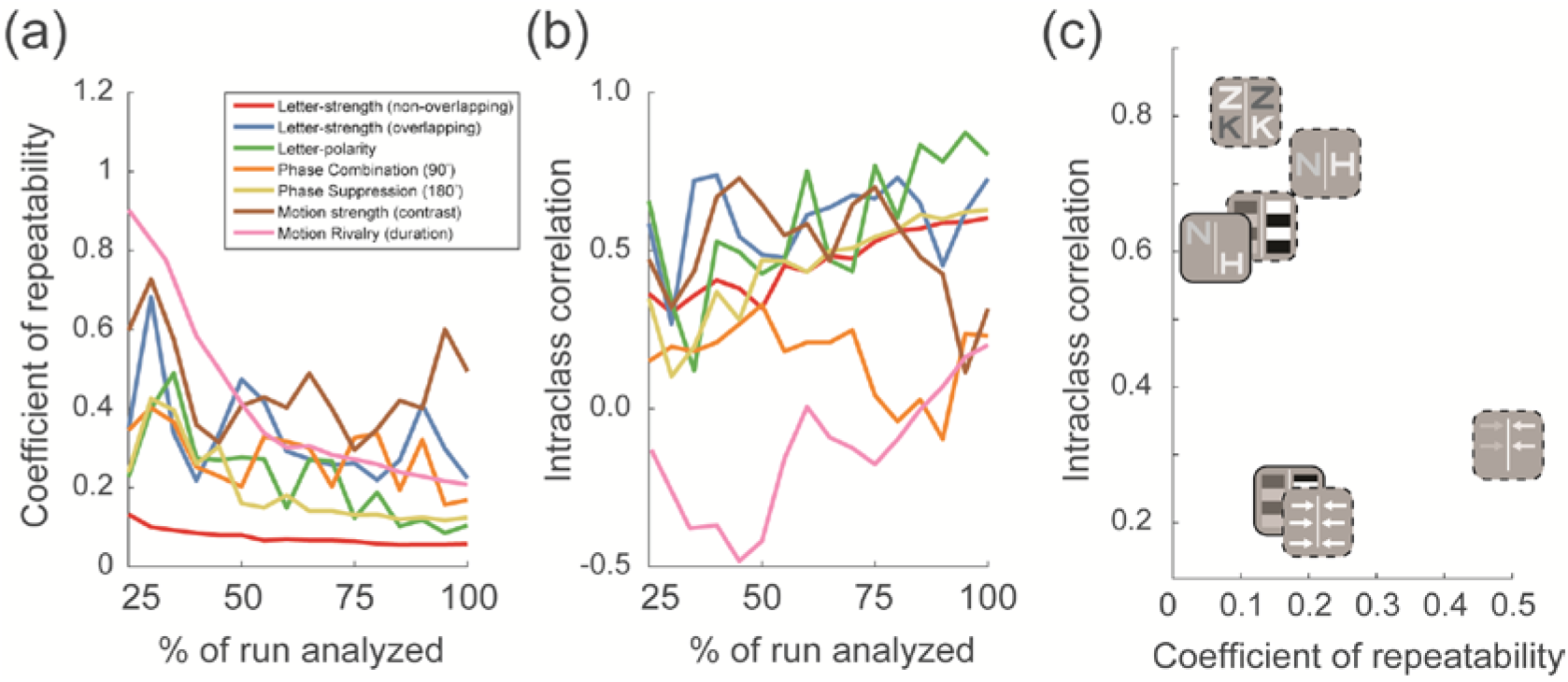
Plot of the evolution of (a) coefficient of repeatability (lower is better) and (b) interclass correlation (higher is better) over run-length (expressed as a percentage of the run analysed, e.g. 100%=20 trials) for the seven balance tests. The tests are listed in legend-panel (a). (c) Plot of intra-class correlation against coefficient of repeatability.

We next assessed whether increased binocular imbalance was associated with poorer stereoacuity and/or with more marked eye-dominance in rivalry. As shown in Figure 5a, we fitted a regression line to the data points corresponding to (y-axis) the magnitude of SED and (x-axis) the stereoacuity estimate from the same participant (mean of the two runs). The mean magnitude of SED was obtained by averaging contrast-BP indexes across tests #1-6 and computing the absolute difference of this value from 0.5. The fit was consistent with only a marginal association between measures (R=0.30), resulting not significant (*p*= 0.10). Figure 5b plots SED magnitude against the mean duration of instances of perceptual dominance, as measured in the rivalry test (#7). These data show a non-significant correlation of R=0.20, *p*=0.30. Participants’ rivalrous percepts lasted for an average of 2.27 seconds (σ=1.42s; median=1.73s).

**Figure 5.**
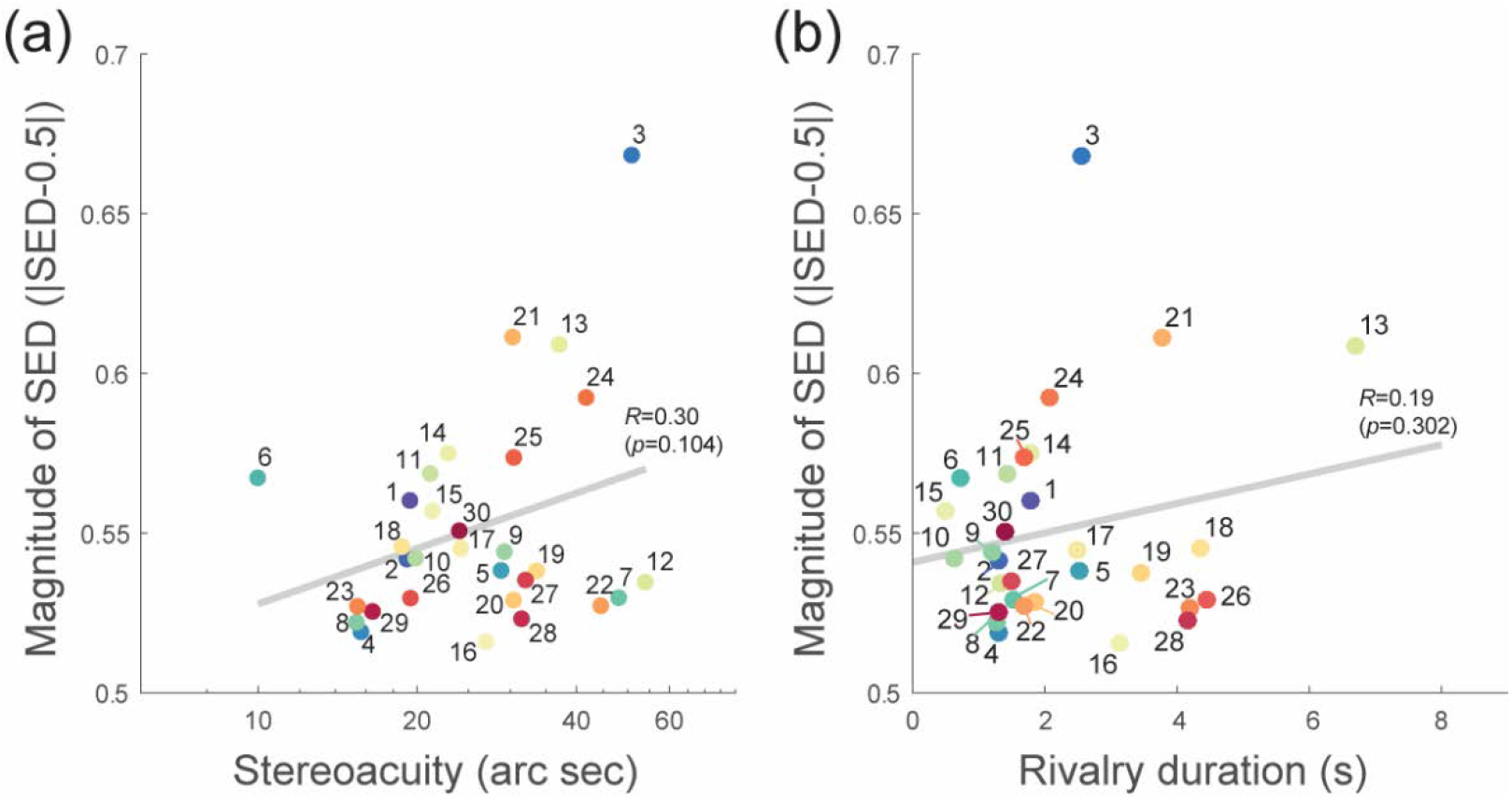
Data from each numbered participant (colour coding of tests is as Figure 3) showing either (a) stereoacuity (task #8) or (b) the duration of perceptual dominance (task #7) plot against magnitude of SED (based on the mean contrast-Balance-Point averaged across test #1-6). All data are averaged across two runs. Magnitude of SED is | SED-0.5|, i.e. 0.50=balanced vision, and 1.0=complete dominance of one eye. On each graph, a regression line has been fit to data, and correlation indexes (*R*) with associated p-values are reported.

## Discussion

We compared eight tests that use dichoptic stimuli to quantify binocular visual function. We measured sensory-eye-dominance (SED) in tests #1-7, and stereoacuity in test #8 in 30 individuals with ostensibly normal vision. All SED tests involved estimation of a *Balance-Point* (BP): an estimate of the extent to which participants relied equally on the two components of a dichoptic stimulus-pair. Specifically, pairs of contrast-modulated stimuli were used in tests #1-6, comprising Sloan letters (adapted from Kwon, Wiecek et al. 2015 -tests #1-2, Bossi, Tailor et al. 2017 -test #3), reciprocally-shifted gratings (from Ding and Sperling 2006 -tests #4-5) or up/down drifting noise-patterns (loosely based on the motion task by Black, Thompson et al. 2011 -test #6). Test #7 (inspired by Dieter, Sy et al. 2016) measured rivalry, as the proportion of dominant-eye instances (presenting the same pairs of stimuli used in #6 but at a fixed contrast). In addition, test #8 measured stereoacuity, using stereo-defined Sloan letters (variant of Frisby, Davis et al. 1996).

We observed a high level of consistency in BPs across tests for each participant. The mean magnitude of SED, obtained by averaging absolute BPs, was 0.55, σ=±0.033, with the majority of participants showing right-eye sensory dominance, in line with previous results (e.g. Ehrenstein, Arnold-Schulz-Gahmen et al. 2005, Pointer 2012, Johansson, Seimyr et al. 2015). Overall, regardless of the statistics used, test #3 showed the best performance in term of validity and reliability; while tests #7 and #4 were the poorest (see Table 2). We found no significant correlation between stereoacuity and SED measures: better binocular balance (i.e. BPs around 0.50) was not associated with better stereoacuity (Figure 5a). Such a null result could simply arise from a lack of variation in stereoacuity amongst our (normally sighted) observers and indeed it has recently been observed that low between-subject variability elicited by robust psychophysical tasks such as stereoacuity, may make them ill-suited for studying individual differences (Hedge, Powell et al. 2017).

*Why are some tests better than others?* In terms of test-validity an ideal test captures a range of BP estimates across participants. We note that binocular imbalance manifests most robustly for stimuli presented at the same location so that estimates of BP from test #1 (non-overlapping letter strength) converge on 0.5. This fundamentally limits the validity of this test for capturing variation in SED within a population of people with normal vision. We would also emphasize that this evaluation is based wholly on a population with ostensibly normal vision; a similar evaluation with a clinical population might yield different results.

In terms of test-reliability, one might expect that tests involving rivalrous stimuli would elicit less reliable responses from participants because, for example, rivalrous percepts typically fluctuate in time. We presented stimuli for relatively short durations, to limit such perceptual alternation. Nevertheless, our expectation - that these stimuli would still elicit more random response – was not borne out by the reliability of tests involving rivalrous stimuli. Rather, our finding of high levels of reliability on these tests suggests that the SED may selectively drive the early stages of a rivalrous percept. This is consistent with earlier reports of participants’ first percept of a rivalrous display predicting their eye dominance (e.g. quantified using percept frequency) calculated for stimuli that alternated over dozens of seconds (Xu, He et al. 2011, Dieter, Sy et al. 2016).

Among the contrast-balance tests, those which rival in motion (#6) and form (#2) show the poorest test-retest reliability (CoR). By contrast, tests #3 and 5, which only rival in polarity, exhibit better reliability. Note (see Figure 4, a-b) that overall this is true throughout a testing session: tests #2 and 6 show a poorer coefficient of reliability compared to most of the other tests and the relative intra-class correlation index gets worse (tests #6) or does not improve (test #2) towards the end of a session. It is possible our results were influenced by our use of a fixed test-order. However if any serial effects (such as adaptation) contributed to our findings (e.g. of poorer performance on tests such as #2 and 6), we would expect results to degrade over the course of a run. This was not the case (as visible in Figure 4a-b). Instead, we believe that the different performance on tests #3 and 5 compared to tests #2 and 6 arises from polarity-rivalry removing the need for participants to explicitly judge relative stimulus strength. For two rivalrous, same-polarity but different-identity targets (e.g. test #2), observers likely judge relative stimulus-strength by comparing two independent estimates of contrast. In this case *binocular imbalance manifests as a difference in magnitude of the two cues* (i.e. one will be “stronger/whiter/brighter” than the other). In contrast, the appearance of two rivalrous, opposite-polarity but same-identity targets (test #3), alternates over time with the *target-pair* appearing as either left-black/right-white or left-white/right-black. In this scenario *binocular imbalance manifests as a difference in probability that one state will dominate over the other*. In other words, for test #2 the observer is faced with a judgement of a subtle difference in perceived luminance, whereas in test #3 they are always judging which of two (apparently) black and white letters is white. This task is straightforward for our adult participants and sufficiently simple for children that we can reliably administer a very similar test using an unsupervised measurement system, as part of a home-based binocular amblyopia therapy (Bossi, Tailor et al. 2017).

*How could new tests fit into clinical practice?* There is an unmet need for accessible, valid and reliable methods to measure binocularity in standard care. In the clinic, binocularity is frequently assessed measuring stereoacuity. However, 1-14% of the population is stereo-blind (Bosten, Goodbourn et al. 2015), and binocular function is necessary but not sufficient for stereoacuity. Moreover, standard stereo-tests (e.g. Frisby, randot) result in non-measurable stereoacuity in cases where compensation for visual deficits (e.g. balancing stimulus visibility across the eyes) allow patients to see stereo (Tytla, Lewis et al. 1993), or achieve binocular summation (Baker, Meese et al. 2007). We therefore would anticipate SED being able to (a) quantify subtle binocular deficits in people with normal stereoacuity and (b) reveal the presence of binocular capacity in the absence of measurable stereoacuity. Our current results speak to (a) – we observe a range of SEDs in our test group – albeit centred on 0.5 or balanced vision. Differences between observers are stable, both in terms of test-retest reliability and in terms of agreement amongst different tests of SED. The current study does not speak to (b) since we have not targeted individuals with abnormal binocular vision. Inclusion of participants with poor binocular vision (the subject of an ongoing study) may influence the outcome of our evaluation. For example, the excellent test-retest reliability of test #1 may outweigh its poor test validity as patients are more likely to exhibit a wider range of larger BPs which may drive this test well but could lead to ceiling effects in more sensitive tests. We further note that patients with abnormal binocular vision will exhibit a wider range of stereoacuities allowing us to more fully explore the relationship between SED and stereoacuity.

Other clinical tests are problematic for their own reasons. For example, tests of “sighting-dominance” (e.g. Miles or Porta tests) deliver binary “left” or “right” measures of eye-dominance. Further, sighting-dominance can be inconsistent, influenced by gaze direction (Khan and Crawford 2001), and by the test used (Rice, Leske et al. 2008). The only continuous estimate of eye-dominance available to clinicians comes from Sbisa bars which rely on the clinician’s judgement of what stimulus level induces a patient to report diplopia. Our procedures have much in common with the Sbisa bar but automate stimulus selection, have the patient perform a forced-choice task, and use all of the response-information to calculate the balance point. This we believe will contribute to superior test reliability and better compliance from patients.

It is important to note that the sight-dominant eye does not necessarily support better visual acuity (Pointer 2007), and laterality - measured with either SED or sight-dominance - only agrees in 50% of cases (Pointer 2012). Further, tests of SED (Suttle, Alexander et al. 2009, Pointer 2012) as well as those of other forms of binocularity, such as sight dominance (Rice, Leske et al. 2008) and stereoacuity (Heron and Lages 2012), produce results that differ depending on the particular visual task and test-conditions. Clearly, both sensory and motor aspects are involved in binocularity. Thus, a complete assessment of functional binocular vision requires more than one type of binocular measurement. The lack of correlation between our (reliable) measures of SED and stereo-acuity suggests these tests are complementary. Future research could more directly focus on determining the sub-processes that support complete binocular vision, and developing a more complete battery of tests to probe them.

*SED and amblyopia* There is increasing interest in the feasibility of treating amblyopia in older children and adults, either with occlusion therapy (Erdem, Cinar et al. 2011, Kishimoto, Fujii et al. 2014) or other monocular (Evans, Yu et al. 2011) and/or binocular methodologies, such as dichoptic training and videogame play (Tsirlin, Colpa et al. 2015). Based on the idea that interocular suppression is a cause not a symptom of amblyopia it has been proposed that binocular treatments achieve their results by reducing interocular suppression. However, a few studies now suggest that therapeutic benefit cannot be underpinned solely by a reduction in interocular suppression (Vedamurthy, Nahum et al. 2015, Bossi, Tailor et al. 2017). There a number of current hypotheses about the mechanism supporting visual improvement but importantly there are also a large number of different measures of SED in use across different laboratories. The assumption that all tests of SED tap the same mechanism may not be true and this may hamper comparative evaluation of the outcomes of different binocular therapies. Having a standard set of tests to measure binocularity could be useful in clarifying how and in which sense interocular suppression, as quantified by the magnitude of imbalance in SED, affects the therapeutic outcome for, and the mechanism of, amblyopia.

## Conclusion

We compared several tests for rapidly quantifying sensory-eye dominance and a test of stereoacuity. A judgement of which of two dichoptically-superimposed opposite contrast-polarity patterns dominated the participant’s percept (test #3) supports a reliable and sensitive measure of sensory-eye dominance in only 20 trials. Practical and reliable measure of SED have application in amblyopia research and as part of a more thorough assessment of binocularity in the clinic.

## Acknowledgements

We thank all the participants and Steven Lim and Young Oh for assistance with data collection. This work was funded by a UCL Impact award to SCD and by a grant from *Cure Kids, NZ* to SCD & LH. SCD is a co-applicant on a US Patent *“Quantification of inter-ocular suppression in binocular vision impairment”* 15/312,376 which may cover one or more of the (letter-based) tests described here.

## References

Babu, R. J., S. Clavagnier, W. R. Bobier, B. Thompson and R. F. Hess (2017). “Regional Extent of Peripheral Suppression in Amblyopia.” Invest Ophthalmol Vis Sci 58(4): 2329–2340.

Baker, D. H., T. S. Meese and R. F. Hess (2008). “Contrast masking in strabismic amblyopia: attenuation, noise, interocular suppression and binocular summation.” Vision Res 48(15): 1625–1640.

Baker, D. H., T. S. Meese, B. Mansouri and R. F. Hess (2007). “Binocular summation of contrast remains intact in strabismic amblyopia.” Invest Ophthalmol Vis Sci 48(11): 5332–5338.

Baker, D. H., T. S. Meese and R. J. Summers (2007). “Psychophysical evidence for two routes to suppression before binocular summation of signals in human vision.” Neuroscience 146(1): 435–448.

Birch, E., C. Williams, J. Drover, V. Fu, C. Cheng, K. Northstone, M. Courage and R. Adams (2008). “Randot Preschool Stereoacuity Test: normative data and validity.” J aapos 12(1): 23–26.

Birch, E. E. (2013). “Amblyopia and binocular vision.” Progress in Retinal and Eye Research 33(0): 67–84.

Birch, E. E., S. L. Li, R. M. Jost, S. E. Morale, A. De La Cruz, D. Stager, Jr., L. Dao and D. R. Stager, Sr., (2015). “Binocular iPad treatment for amblyopia in preschool children.” J AAPOS 19(1): 6–11.

Black, J. M., R. F. Hess, J. R. Cooperstock, L. To and B. Thompson (2012). “The measurement and treatment of suppression in amblyopia.” J Vis Exp(70): e3927.

Black, J. M., B. Thompson, G. Maehara and R. F. Hess (2011). “A compact clinical instrument for quantifying suppression.” Optom Vis Sci 88(2): E334–343.

Bossi, M., V. K. Tailor, E. J. Anderson, P. J. Bex, J. A. Greenwood, A. Dahlmann-Noor and S. C. Dakin (2017). “Binocular Therapy for Childhood Amblyopia Improves Vision Without Breaking Interocular Suppression.” Invest Ophthalmol Vis Sci 58(7): 3031–3043.

Bosten, J. M., P. T. Goodbourn, A. J. Lawrance-Owen, G. Bargary, R. E. Hogg and J. D. Mollon (2015). “A population study of binocular function.” Vision Res 110(Pt A): 34–50.

Carkeet, A. (2015). “Exact parametric confidence intervals for Bland-Altman limits of agreement.” Optom Vis Sci 92(3): e71–80.

Cotter, S. A., L. A. Cyert, J. M. Miller and G. E. Quinn (2015). “Vision screening for children 36 to <72 months: recommended practices.” Optom Vis Sci 92(1): 6–16.

Crawford, L. C. J. and H. J. Griffiths (2015). “The repeatability of the Sbisa bar for testing density of suppression.” British and Irish Orthoptic Journal 12: 35–40.

Cumming, B. G. and G. C. DeAngelis (2001). “The physiology of stereopsis.” Annu Rev Neurosci 24: 203–238.

Dieter, K. C., J. L. Sy and R. Blake (2016). “Individual differences in sensory eye dominance reflected in the dynamics of binocular rivalry.” Vision Res.

Ding, J. and G. Sperling (2006). “A gain-control theory of binocular combination.” Proceedings of the National Academy of Sciences of the United States of America 103(4): 1141–1146.

Dorr, M., T. Elze, H. Wang, Z. L. Lu, P. J. Bex and L. Lesmes (2017). New precision metrics for contrast sensitivity testing.

Ehrenstein, W. H., B. E. Arnold-Schulz-Gahmen and W. Jaschinski (2005). “Eye preference within the context of binocular functions.” Graefe‘s Archive for Clinical and Experimental Ophthalmology 243(9): 926–932.

Erdem, E., G. Y. Cinar, D. Somer, N. Demir, A. Burcu and F. Ornek (2011). “Eye patching as a treatment for amblyopia in children aged 10-16 years.” Jpn J Ophthalmol 55(4): 389–395.

Evans, B. J. W., C. S. Yu, E. Massa and J. E. Mathews (2011). “Randomised controlled trial of intermittent photopic stimulation for treating amblyopia in older children and adults.” Ophthalmic Physiol Opt 31: 56–68.

Ferris, F. L., 3rd, A. Kassoff, G. H. Bresnick and I. Bailey (1982). “New visual acuity charts for clinical research.” Am J Ophthalmol 94(1): 91–96.

Fricke, T. R. and J. Siderov (1997). “Stereopsis, stereotests, and their relation to vision screening and clinical practice.” Clinical and Experimental Optometry 80(5): 165–172.

Frisby, J. P., H. Davis and K. McMorrow (1996). “An improved training procedure as a precursor to testing young children with the Frisby Stereotest.” Eye (Lond) 10 (Pt 2): 286–290.

Hamm, L., Z. Chen, J. Li, J. Black, S. Dai, J. Yuan, M. Yu and B. Thompson (2017). “Interocular suppression in children with deprivation amblyopia.” Vision Res 133: 112–120.

Hedge, C., G. Powell and P. Sumner (2017). “The reliability paradox: Why robust cognitive tasks do not produce reliable individual differences.” Behav Res Methods.

Hess, R. F. and B. Thompson (2015). “Amblyopia and the binocular approach to its therapy.” Vision Res.

Huang, C.-B., J. Zhou, Y. Zhou and Z.-L. Lu (2010). “Contrast and phase combination in binocular vision.” PLoS One 5(12): e15075.

Huang, C. B., J. Zhou, Z. L. Lu, L. Feng and Y. Zhou (2009). “Binocular combination in anisometropic amblyopia.” J Vis 9(3): 17 11–16.

Johansson, J., G. Ö. Seimyr and T. Pansell (2015). “Eye dominance in binocular viewing conditions.” Journal of vision 15(9): 21–21.

Julesz, B. (1971). Foundations of Cyclopean Perception, University of Chicago Press.

Kehrein, S., T. Kohnen and M. Fronius (2016). “Dynamics of Interocular Suppression in Amblyopic Children during Electronically Monitored Occlusion Therapy: First Insight.” Strabismus 24(2): 51–62.

Khan, A. Z. and J. D. Crawford (2001). “Ocular dominance reverses as a function of horizontal gaze angle.” Vision Research 41(14): 1743–1748.

Kishimoto, F., C. Fujii, Y. Shira, K. Hasebe, I. Hamasaki and H. Ohtsuki (2014). “Outcome of conventional treatment for adult amblyopia.” Jpn J Ophthalmol 58(1): 26–32.

Kwon, M., Z. L. Lu, A. Miller, M. Kazlas, D. G. Hunter and P. J. Bex (2014). “Assessing binocular interaction in amblyopia and its clinical feasibility.” PLoS One 9(6): e100156.

Kwon, M., E. Wiecek, S. C. Dakin and P. J. Bex (2015). “Spatial-frequency dependent binocular imbalance in amblyopia.” Sci Rep 5: 17181.

Li, J., R. F. Hess, L. Y. Chan, D. Deng, X. Chen, M. Yu and B. S. Thompson (2013). “How best to assess suppression in patients with high anisometropia.” Optom Vis Sci 90(2): e47–52.

Li, J., R. F. Hess, L. Y. Chan, D. Deng, X. Yang, X. Chen, M. Yu and B. Thompson (2013). “Quantitative measurement of interocular suppression in anisometropic amblyopia: a case-control study.” Ophthalmology 120(8): 1672–1680.

Li, J., C. S. Lam, M. Yu, R. F. Hess, L. Y. Chan, G. Maehara, G. C. Woo and B. Thompson (2010). “Quantifying sensory eye dominance in the normal visual system: a new technique and insights into variation across traditional tests.” Invest Ophthalmol Vis Sci 51(12): 6875–6881.

Li, J., B. Thompson, D. Deng, L. Y. Chan, M. Yu and R. F. Hess (2013). “Dichoptic training enables the adult amblyopic brain to learn.” Curr Biol 23(8): R308–309.

Mansouri, B., B. Thompson and R. F. Hess (2008). “Measurement of suprathreshold binocular interactions in amblyopia.” Vision Res 48(28): 2775–2784.

McCormick, A., R. Bhola, L. Brown, D. Squirrel, J. Giles and I. Pepper (2002). “Quantifying relative afferent pupillary defects using a Sbisa bar.” Br J Ophthalmol 86(9): 985–987.

McGraw, K. O. and S. P. Wong (1996). “Forming inferences about some intraclass correlation coefficients.” Psychological methods 1(1): 30.

McKee, S. P., D. M. Levi and J. A. Movshon (2003). “The pattern of visual deficit in amblyopia.” Journal of Vision 3(5): 380–405.

Miles, W. R. (1930). “Ocular Dominance in Human Adults.” The Journal of General Psychology 3(3): 412–430.

O‘Connor, A. R., E. E. Birch, S. Anderson and H. Draper (2010). “The functional significance of stereopsis.” Invest Ophthalmol Vis Sci 51(4): 2019–2023.

Ooi, T. L. and Z. J. He (2001). “Sensory eye dominance.” Optometry 72(3): 168–178.

Ooi, T. L., Y. R. Su, D. M. Natale and Z. J. He (2013). “A push-pull treatment for strengthening the ‘lazy eye’ in amblyopia.” Curr Biol 23(8): R309–310.

Piano, M. and D. Newsham (2015). “A pilot study examining density of suppression measurement in strabismus.” Strabismus 23(1): 14–21.

Pointer, J. S. (2012). “Sighting versus sensory ocular dominance.” Journal of Optometry 5(2): 52–55.

Prins, N. and F. A. A. Kingdom. (2009). “Palamedes: Matlab routines for analyzing psychophysical data.”, from http://www.palamedestoolbox.org

Rice, M. L., D. A. Leske, C. E. Smestad and J. M. Holmes (2008). “Results of Ocular Dominance Testing Depend on Assessment Method.” Journal of AAPOS : the official publication of the American Association for Pediatric Ophthalmology and Strabismus / American Association for Pediatric Ophthalmology and Strabismus 12(4): 365–369.

Tailor, V., S. Balduzzi, S. Hull, J. Rahi, C. Schmucker, G. Virgili and A. Dahlmann-Noor (2014). “Tests for detecting strabismus in children age 1 to 6 years in the community.” Cochrane Database of Systematic Reviews(7).

Tsirlin, I., L. Colpa, H. C. Goltz and A. M. Wong (2015). “Behavioral Training as New Treatment for Adult Amblyopia: A Meta-Analysis and Systematic Review.” Invest Ophthalmol Vis Sci 56(6): 4061–4075.

Tytla, M. E., T. L. Lewis, D. Maurer and H. P. Brent (1993). “Stereopsis after congenital cataract.” Invest Ophthalmol Vis Sci 34(5): 1767–1773.

Vaz, S., T. Falkmer, A. E. Passmore, R. Parsons and P. Andreou (2013). “The Case for Using the Repeatability Coefficient When Calculating Test–Retest Reliability.” PLoS ONE 8(9): e73990.

Vedamurthy, I., M. Nahum, D. Bavelier and D. M. Levi (2015). “Mechanisms of recovery of visual function in adult amblyopia through a tailored action video game.” Sci Rep 5: 8482.

Vedamurthy, I., M. Nahum, S. J. Huang, F. Zheng, J. Bayliss, D. Bavelier and D. M. Levi (2015). “A dichoptic custom-made action video game as a treatment for adult amblyopia.” Vision Res 114: 173–187.

Watson, A. B. and D. G. Pelli (1983). “QUEST: a Bayesian adaptive psychometric method.” Percept Psychophys 33(2): 113–120.

Wheatstone, C. (1838). “Contributions to the Physiology of Vision.--Part the First. On Some Remarkable, and Hitherto Unobserved, Phenomena of Binocular Vision.” Philosophical transactions of the Royal Society of London 128: 371–394.

Wiecek, E., K. Lashkari, S. C. Dakin and P. Bex (2015). “Metamorphopsia and interocular suppression in monocular and binocular maculopathy.” Acta Ophthalmol 93(4): e318–320.

Wong, A. M. F. (2012). “New concepts concerning the neural mechanism of amblyopia and their clinical implications.” Can J Ophthalmol 47(5): 399–409.

Xu, J. P., Z. J. He and T. L. Ooi (2011). “A binocular perimetry study of the causes and implications of sensory eye dominance.” Vision research 51(23-24): 2386–2397.

Zhou, J., S. Clavagnier and R. F. Hess (2013). “Short-term monocular deprivation strengthens the patched eye‘s contribution to binocular combination.” J Vis 13(5).

Zhou, J., P. C. Huang and R. F. Hess (2013). “Interocular suppression in amblyopia for global orientation processing.” J Vis 13(5): 19.

Zhou, J., A. Reynaud and R. F. Hess (2014). “Real-time modulation of perceptual eye dominance in humans.” Proc Biol Sci 281(1795).

Zhou, J., B. Thompson and R. F. Hess (2013). “A new form of rapid binocular plasticity in adult with amblyopia.” Sci Rep 3: 2638.

